# Rapid and Cost-Effective Digital Quantification of RNA Editing and Maturation in Organelle Transcripts

**DOI:** 10.1101/2025.06.23.661184

**Authors:** Zhihua Hua

**Author notes:** **Corresponding author:** Zhihua Hua, Department of Environmental and Plant Biology, Ohio University, Athens, Ohio 45701 USA.

## Abstract

RNA editing and maturation are critical regulatory mechanisms in plant organelles, yet their quantification remains technically challenging. Traditional Sanger sequencing lacks sensitivity and reproducibility, whereas advanced next-generation sequencing (NGS) approaches, such as rRNA-depleted RNA-seq or targeted amplicon-seq, involve high costs, complex workflows, and limited accessibility. To address these limitations, we developed a rapid and cost-effective long-read sequencing approach, termed premium PCR sequencing, for digital quantification of RNA-editing and intron retention events in targeted chloroplast transcripts. This method combines multiplexed high-fidelity PCR amplification with Oxford Nanopore sequencing and custom in-house Perl and Python scripts for streamlined data processing, including barcode-based demultiplexing, strand reorientation, alignment to a pseudo-genome, manual editing-site inspection, and splicing variant identification and comparison. Using this platform, we analyzed the *ndhB* and *ndhD* transcripts, two chloroplast *NAD(P)H dehydrogenase* genes with the highest number of known editing sites, in an inducible CRISPR interference (iCRISPRi) system targeting *MORF2*, a key RNA-editing factor. Our results revealed MORF2 dosage-dependent reductions in C-to-U editing efficiency, with significant defects observed in the strongly repressed *P1-12* line. Moreover, we identified an accumulation of intron-retaining *ndhB* transcripts, specifically in Dex-treated iCRISPRi lines, indicating impaired chloroplast splicing functions upon *MORF2* suppression. The platform achieves single-molecule resolution, robust reproducibility, and high read coverage across biological replicates at a fraction of the cost of lncRNA-seq, making it broadly accessible. This study establishes premium PCR sequencing as a versatile, scalable, and affordable tool for targeted post-transcriptional analysis in plant organelles and expands our understanding of MORF2’s role in chloroplast RNA maturation.

**Significance Statement:** We present a rapid, affordable, and reproducible method for digital quantification of RNA editing and intron retention in plant organellar transcripts using nanopore-based long-read sequencing. This platform overcomes key limitations of existing approaches and enables routine, site-specific analysis of post-transcriptional regulation in organelles, including RNA editing and splicing, making it broadly accessible to researchers studying plastid biology, stress responses, and organelle–nucleus communication.

## Introduction

The presence of multiple genomes in eukaryotic cells necessitates distinct regulatory mechanisms to coordinate gene expression between nuclear and organellar genomes, such as those found in mitochondria and chloroplasts in plant cells. This genomic compartmentalization has driven the evolution of intricate signaling networks essential for proper development and physiological adaptations through coordinated gene expression (Woodson and Chory, 2008, Jarvis and Lopez-Juez, 2013). Among these regulatory mechanisms, RNA editing represents a distinctive post-transcriptional modification predominantly observed in mitochondria and chloroplasts. Through site-specific conversion primarily of cytidine (C) to uridine (U), RNA editing modifies transcripts to restore conserved codons, generates functional AUG start codons, eliminates premature stop codons, and influences RNA structure, splicing, and stability (Takenaka *et al*., 2013, Small *et al*., 2020). These modifications are critical for organelle biogenesis and have been increasingly recognized as dynamic regulators of stress responses and developmental signaling (Hao *et al*., 2021, Hu *et al*., 2024, Mohamed *et al*., 2024).

RNA editing is particularly abundant in the organellar transcripts of land plants, with approximately 40 and 500 editing sites identified in chloroplast and mitochondrial RNAs, respectively, in *Arabidopsis thaliana* (Chateigner-Boutin and Small, 2007, Bentolila *et al*., 2008). These editing events often lead to amino acid substitutions crucial for the functionality of proteins involved in photosynthetic electron transport, ATP synthesis, and ribosome assembly (Sun *et al*., 2013, Takenaka *et al*., 2013, Small *et al*., 2020). Recent studies further demonstrated dynamic regulation of editing efficiency in response to environmental stimuli such as temperature (Cui *et al*., 2019, Wu *et al*., 2025), light (Hu *et al*., 2025), and drought (Mohamed *et al*., 2024), underscoring RNA editing as an adaptable mechanism for environmental acclimation and signaling.

Despite its biological importance, quantitative analysis of RNA-editing efficiency remains technically challenging. Multiple methodological strategies have been employed to measure RNA-editing efficiency in plant organelles, each offering distinct advantages and limitations in terms of resolution, scalability, cost, and accessibility.

The traditional approach, Sanger sequencing of reverse transcription PCR (RT-PCR) products, remains widely used for small-scale validation of editing events (Takenaka *et al*., 2012, Zhao *et al*., 2019, Tang *et al*., 2024, Wu *et al*., 2025). This method provides direct visualization of cytosine (C) to thymine (T) peak shifts in sequencing chromatograms and requires only basic laboratory infrastructure (Takenaka *et al*., 2012). However, it is inherently low-throughput, semi-quantitative, and exhibits poor reproducibility, particularly when editing efficiencies are modest (<20%) (Tang *et al*., 2024). Consequently, Sanger sequencing is not suitable for extensive comparative analyses across multiple genes, conditions, or biological replicates.

To improve throughput and quantification, strand- and transcript-specific PCR sequencing (STS-PCRseq) and targeted amplicon-seq have been employed. These techniques involve the PCR amplification of specific organelle transcript fragments, followed by the preparation of a next-generation sequencing (NGS) library and deep sequencing on Illumina platforms. STS-PCRseq offers single-nucleotide resolution and high sensitivity, facilitating comprehensive and high-resolution profiling of RNA editing events in plastids and mitochondria of *A. thaliana* (Bentolila *et al*., 2013). However, these targeted approaches still require comprehensive NGS workflows, including PCR amplicon fragmentation, adapter ligation, quality controls, and multiplexed sequencing, which collectively result in high experimental costs, extended turnaround times, and multiple labor-intensive steps. Additionally, they depend heavily on access to sequencing facilities and PCR reproducibility.

In contrast, transcriptome-wide RNA sequencing (RNA-seq) has become increasingly popular for concurrently analyzing RNA editing, gene expression, and splicing, employing specialized bioinformatics pipelines such as ChloroSeq and REDItools (Malbert *et al*., 2018, Lo Giudice *et al*., 2020). However, conventional RNA-seq typically utilizes poly(A) enrichment, preferentially capturing nuclear-encoded transcripts while excluding non-polyadenylated organellar RNAs. This underrepresentation severely limits statistical power for detecting RNA-editing variations (Yapa *et al*., 2023).

To address this limitation, specialized rRNA-depleted total RNA sequencing (lncRNA-seq) protocols have been developed. These protocols employ ribosomal RNA depletion followed by reverse transcription using random primers, facilitating comprehensive and unbiased recovery of organellar transcripts (Xu *et al*., 2023, Tang *et al*., 2024). Although lncRNA-seq significantly improves organellar transcriptome coverage, it remains the most costly method due to both the high price of rRNA depletion kits and the deep sequencing required for sufficient coverage of editing sites.

Collectively, the biological importance of organelle RNA editing and the limitations of current analytical methods underscore the critical need for a rapid, affordable, and scalable approach that can deliver absolute, digital quantification of RNA-editing events at defined sites without the high costs associated with extensive NGS pipelines. Here, we introduce Premium PCR sequencing, a novel method specifically designed for fast, cost-effective, and digital quantification of RNA-editing efficiency at targeted organellar transcript sites, employing Oxford Nanopore’s long-read sequencing platform to capture full-length amplicons.

Premium PCR sequencing fills a critical methodological gap by enabling precise, absolute quantification of RNA-editing events with exceptional sensitivity, rapid turnaround, robust reproducibility, and scalable throughput, while eliminating the need for complex bioinformatics analyses and costly deep sequencing workflows.

## Results

### Transcriptome complexity leads to differential RT-PCR products of chloroplast genes

In one of our previous studies, we developed an inducible CRISPR interference (iCRISPRi) strategy for repressing the expression of *MULTIPLE ORGANELLAR RNA EDITING FACTOR 2* (*MORF2*) by co-expressing a single guide RNA (sgRNA) targeted to the transcriptional start site (TSS) of *MORF2* and a Krüppel-associated-box (KRAB) domain fused to the carboxyl terminus of endonuclease-deactivated Cas9 (dCas9-KRAB). Because *morf2* null mutants are albino and developmentally lethal (Bisanz *et al*., 2003, Takenaka *et al*., 2012, Yapa *et al*., 2023), the expression of *dCas9-KRAB* was driven by a dexamethasone (Dex)-inducible promoter (Aoyama and Chua, 1997), whereas the sgRNA was constitutively expressed under the *Arabidopsis U6-26* promoter (Xing *et al*., 2014). Thus, only upon Dex treatment is dCas9-KRAB expressed and guided by the sgRNA to the TSS of MORF2, enabling its transcriptional repression in a dose-dependent manner. This inducible suppression allows the otherwise lethal *iCRISPRi-MORF2* transgenic plants to remain viable (Yapa *et al*., 2023).

Given the known role of MORF2 in regulating chloroplast RNA-editing process, our previous work also utilized both Sanger sequencing of RT-PCR products and high-throughput short-read RNA-Seq to verify reduced RNA-editing efficiency in Dex-treated, 7-day(d)-old *iCRISPRi-MORF2* seedlings. However, Sanger sequencing lacked sensitivity and reproducibility, while RNA-Seq captured few organellar reads, making quantification inefficient and costly (Yapa *et al*., 2023).

In this study, we revisited the same *iCRISPRi-MORF2* lines to develop a nanopore-based long-read sequencing platform paired with custom bioinformatics tools for accurate and cost-effective analysis of chloroplast RNA editing and transcript maturation (Fig. 1). We focused on *ndhB* and *ndhD*, two chloroplast *NAD(P)H dehydrogenase* genes that harbor the highest number of C-to-U editing sites among chloroplast transcripts (nine and five, respectively; (Bentolila *et al*., 2013)). Notably, *ndhB* also contains a group II intron (Ostheimer *et al*., 2003), enabling additional investigation of MORF2’s potential role in RNA maturation and splicing, given its known interactions with multiple Pentatricopeptide Repeat (PPR) proteins that are implicated in chloroplast RNA splicing (Takenaka *et al*., 2012, Bayer-Csaszar *et al*., 2017, Small *et al*., 2023, Zhang *et al*., 2023) and in maturation of the plastid ribosomal RNAs (Bisanz *et al*., 2003).

**Fig. 1.**
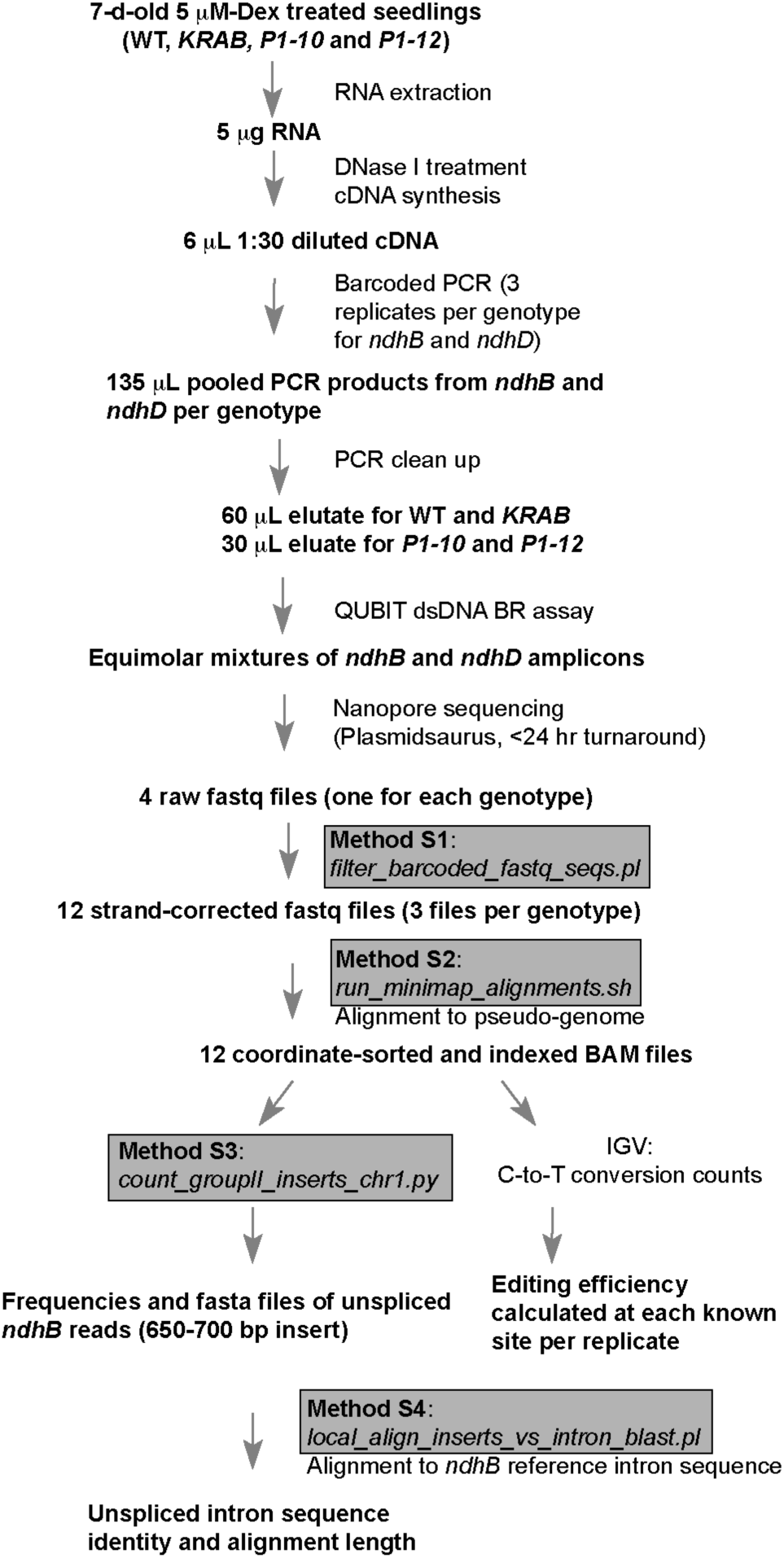
Workflow of premium PCR sequencing and data analysis for RNA-editing efficiency and intron retention. A schematic diagram illustrating the experimental and analytical steps of the premium PCR sequencing platform. Custom open-source bioinformatics scripts (Methods S1– S4) were developed in-house to streamline the workflow for barcode demultiplexing, strand correction, pseudo-genome alignment, editing-site quantification, and intron-retention analysis.

Using quantitative PCR (qPCR), we verified that *MORF2* transcript levels were substantially reduced by 5 μM Dex to 50.2±10.8% in *P1-10* and 20.1±3.9% in *P1-12* relative to WT (mean ± SD). As a control, *KRAB* plants expressing only Dex-inducible *dCas9-KRAB* retained 84.3 ± 3.9% of WT *MORF2* transcript levels (Fig. 2a). This gradient of *MORF2* repression enabled quantitative comparison of its dose-dependent effect on *ndhB* and *ndhD* transcripts.

**Fig. 2.**
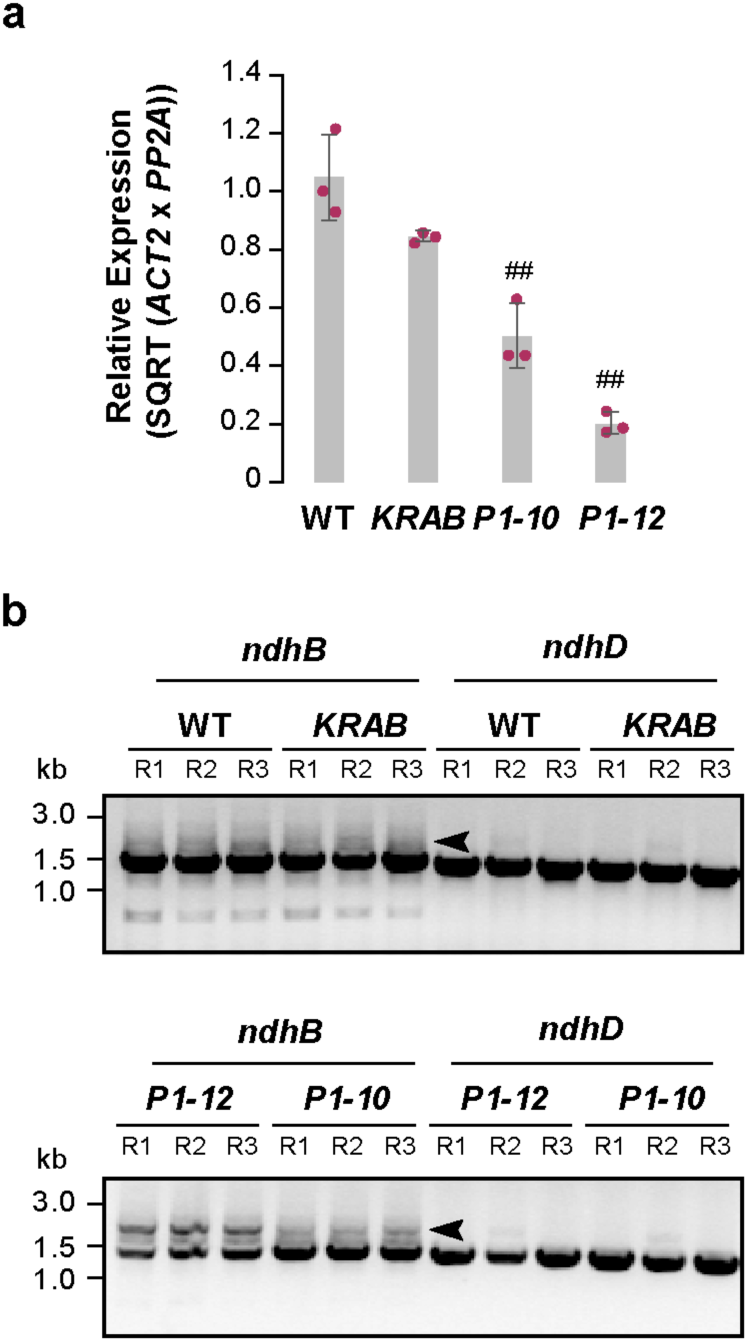
MORF2 suppression by iCRISPRi alters *ndhB* transcript profiles. a) Quantitative PCR analysis showing suppression of *MORF2* transcript levels in 7-day-old seedlings treated with 5 μM Dex. Expression values are normalized to one WT replicate. *ACT2* and *PP2A* were used as internal reference genes. Bars represent the mean ± SD of three biological replicates, each measured in technical duplicate. Individual data points (maroon dots) show biological replicates. # and ## denote statistically significant differences from WT (##P < 0.01; Student’s *t*-test, one-tailed with unequal variance). b) Gel images of RT-PCR products of *ndhB* and *ndhD* from the same cDNAs as used in a). R1-R3 indicate three biological replicates. Forward primers were barcoded for multiplexing. Five microliters of each 50 μL PCR reaction were loaded on a 1% agarose gel stained with ethidium bromide.

To obtain full-length cDNA products for nanopore long-read sequencing, we synthesized first-strand cDNA using a combination of oligo(dT)25 and random hexamer primers, followed by high-fidelity PCR amplification of *ndhB* and *ndhD* cDNAs. The resulting amplicons displayed genotype-dependent differences in both yield and complexity. Robust PCR products were observed for both *ndhB* and *ndhD* in WT and *KRAB*. In contrast, *ndhB* amplicons were moderately and strongly reduced in *P1-10* and *P1-12*, respectively, whereas *ndhD* products were only mildly affected in the same two genotypes, suggesting gene-specific sensitivity of transcript processing to MORF2 suppression (Fig. 2b).

Notably, in addition to the expected full-length *ndhB* band (∼1.4 kb), larger amplicons appeared specifically in Dex-treated *P1-10* and *P1-12* lines. These upper bands correspond to intron-retaining, premature mRNA isoforms (arrowheads, Fig. 2b). These complex banding patterns precluded standard Sanger sequencing for editing analysis, as clean chromatograms could only be obtained from single, specific PCR products. In our previous studies, we had to design internal primers to amplify smaller *ndhB* fragments, followed by gel extraction; however, this process was time-consuming, poorly reproducible, and resulted in inconsistent sequencing results (Yapa *et al*., 2023).

### High read coverage of *ndhB* and *ndhD* transcripts from nanopore long-read sequencing

Since nanopore long-read sequencing analyzes DNA strands at the single-molecule level, we barcoded the *ndhB* and *ndhD* PCR amplicons from each biological replicate using their forward primers and pooled the three replicates, 135 μL in total (each with 45 μL per replicate from an initial 50 μL PCR reaction), without performing an additional gel extraction step. Based on the stronger band intensity of *ndhB* and *ndhD* PCR products observed in WT and *KRAB* than in *P1-10* and *P1-12* from DNA gel electrophoresis (Fig. 1b), we eluted the cleaned DNA in 60 μL for WT and *KRAB*, and 30 μL for *P1-10* and *P1-12*. Using the Qubit double-stranded (ds)DNA Broad Range (BR) assay, we confirmed that each cleaned product exceeded 50 ng/μL in concentration.

To ensure an unbiased representation of *ndhB* and *ndhD* PCR amplicons in each final nanopore sequencing library, we prepared equimolar mixtures of *ndhB* and *ndhD* PCR products from each genotype (Table 1). As expected, the resulting sequencing output yielded comparable read counts for ndhB and ndhD within each genotype, with totals ranging from 891 to 1300 reads (Table S1; Appendix S1).

**Table 1.**
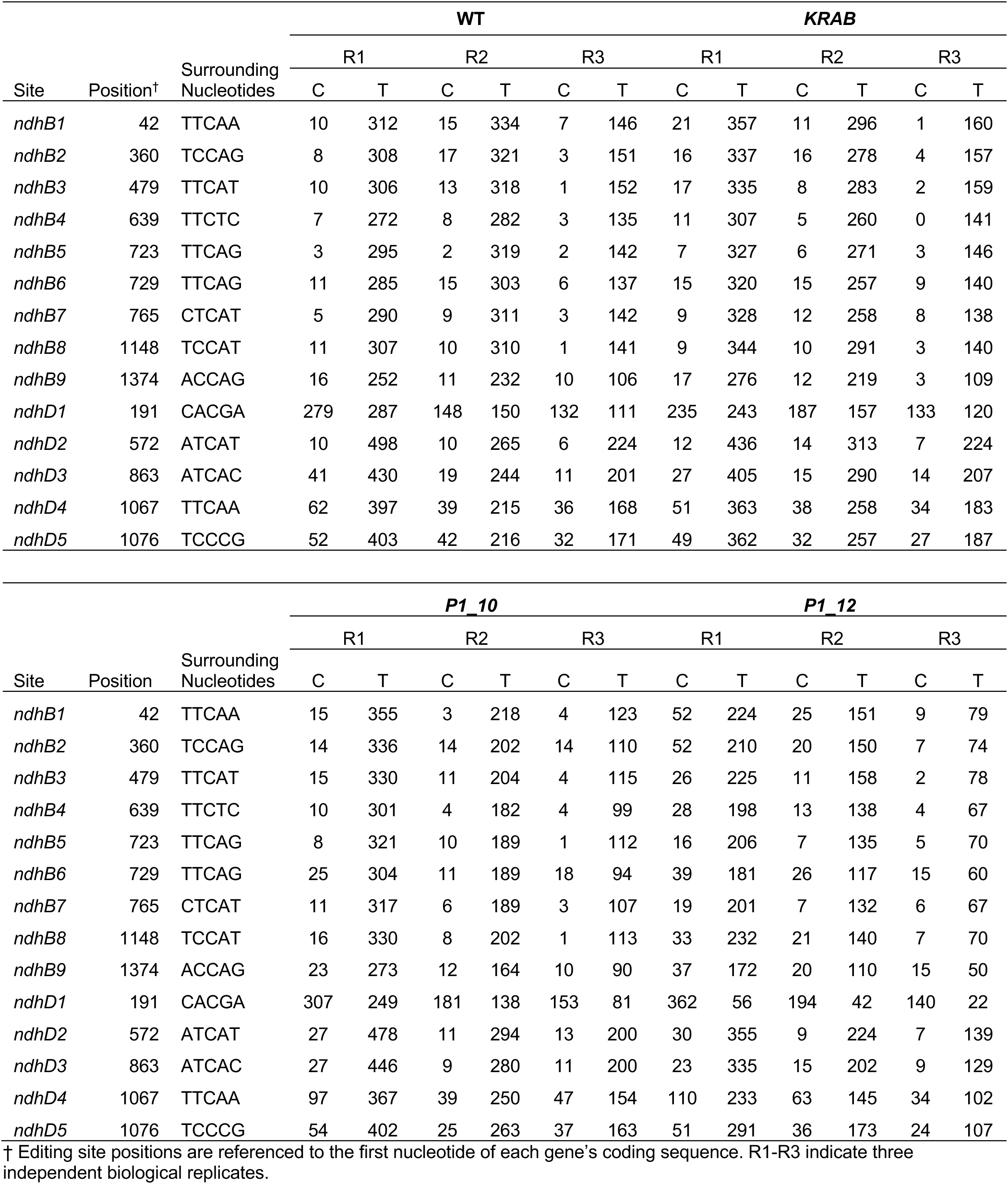
Counts of cytidine (C) and thymine (T) nucleotides at each known RNA-editing site in *ndhB* and *ndhD* RT-PCR products from four genotypes.

Interestingly, the barcode sequence appears to influence the number of reads recovered from each sample. Among the three barcode samples, ATGCTAGC (Replicate 1) produced the highest number of reads, followed by CGTACGTA (Replicate 2) and TACGATCG (Replicate 3) (Table S2), suggesting barcode sequence may affect read recovery efficiency in nanopore sequencing.

### Digital quantification confirmed MORF2 dose-dependent RNA-editing changes

The high read coverage from each barcoded PCR sequencing result, ranging from 122 to 636 (Table S2), provided strong statistical power to digitally quantify RNA-editing efficiency across genotypes. Using an in-house developed Perl script that incorporates three BioPerl modules — *Bio::SeqIO*, *Bio::Seq*, and *Bio::Seq::Quality* (Fig. 1; Method S1) —we processed the sequence and quality data in each raw FASTQ file (Appendix S1) into strand-corrected versions. Each read sequence and corresponding quality scores were oriented to match the sense strand of *ndhB* and *ndhD*. Based on the presence of a specific barcode within each sequence, we generated three strand-corrected FASTQ files per raw input file for the *ndhB* and *ndhD* amplicons, each representing the sequencing result of three biological replicates from each genotype, as indicated in Fig. 2b.

We then aligned the strand-corrected reads to a pseudo-genome (Fig. 1; Method S2; Appendix S2), which is composed of six synthetic chromosomes, each representing a barcoded version of the *ndhB* or *ndhD* reference cDNA sequences. This enabled us to generate three Binary Alignment/Map (BAM) and BAM index (BAI) files corresponding to the three biological replicates of each genotype. Since the positions of the known editing sites were predefined, we manually quantified the number of cytidine (C) and thymine (T) nucleotides at each site across genotypes by visualizing the indexed BAM files using the Integrative Genomics Viewer (IGV) browser (Table 1).

In total, we observed 65 to 556 valid reads per editing site across all sequencing files (median count = 217), enabling robust C-to-U editing efficiency calculations (Table 1). Our results revealed statistically significant reductions in editing efficiency at six out of nine *ndhB* and two out of five *ndhD* editing sites in *P1-12* compared to WT (Fig. 3; *P* < 0.05, Student’s t-test, two-tailed with unequal variance). In contrast, no statistically significant differences were found in *P1-10*, suggesting that RNA-editing defects, at least in *ndhB* and *ndhD,* occur in a MORF2 dose-dependent manner. These findings are consistent with our prior Sanger sequencing results at five *ndhB* editing sites (Yapa *et al*., 2023).

**Fig. 3.**
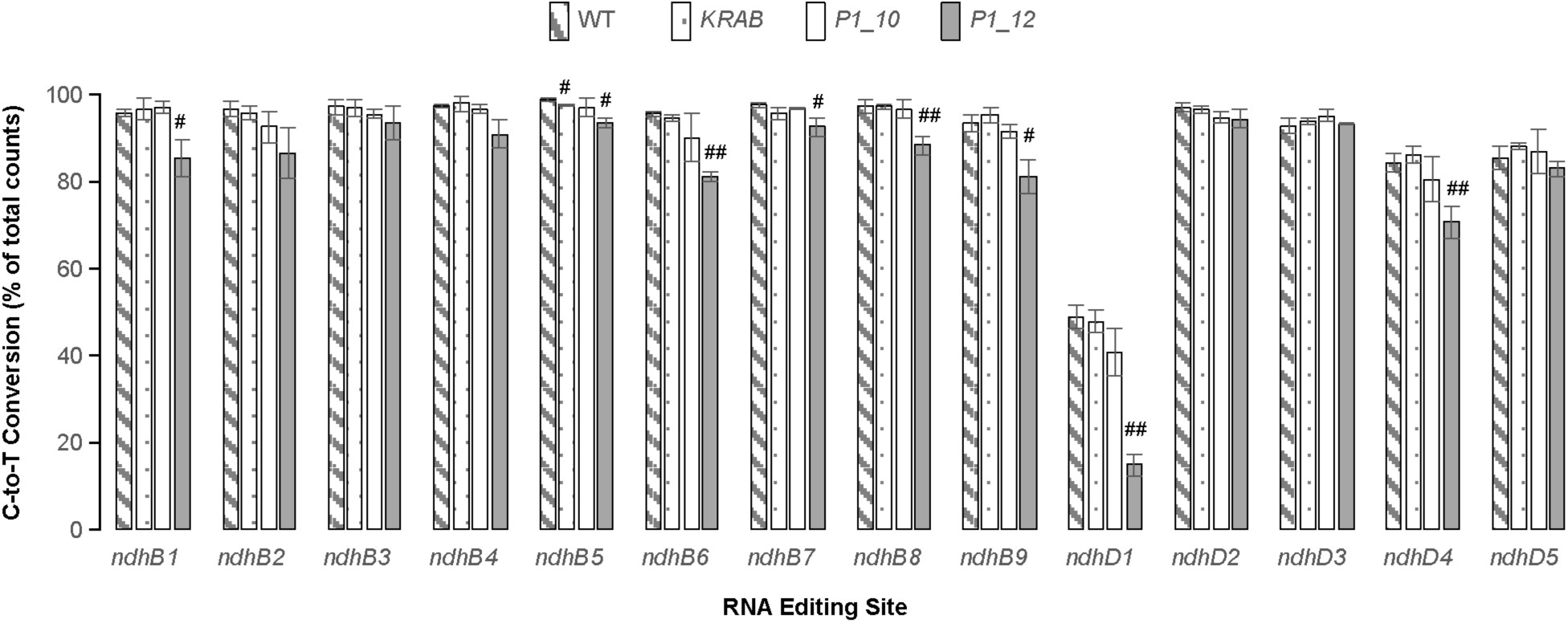
Reduced *MORF2* transcription impairs RNA editing in *ndhB* and *ndhD* transcripts. C-to-U RNA-editing efficiencies of *ndhB* and *ndhD* were quantified from nanopore long-read sequencing of multiplexed RT-PCR amplicons, as shown in Fig. 2b. Data represent the mean ± SD of three biological replicates (see Table 1). Refer to Table 1 for the positions and surrounding nucleotide contexts of each RNA-editing site. Statistically significant differences from WT are indicated (#P < 0.05, ##P < 0.01; Student’s *t*-test, two-tailed with unequal variance).

Interestingly, we also detected a modest but statistically significant reduction in editing efficiency at one *ndhB* editing site (position 723, relative to the first nucleotide in the coding sequence) in *KRAB* compared to WT (99.0% in WT vs. 97.9% in *KRAB*; *P* = 0.035, Student’s t-test, two-tailed with unequal variance), suggesting a possible off-target or stress-related effect of dCas9-KRAB expression.

### A high frequency of unprocessed premature *ndhB* transcripts in *P1-10* and *P1-12* upon Dex treatment

The presence of an additional PCR product band with a molecular weight higher than the expected 1,405 bp *ndhB* transcript (Fig. 2b, indicated with an arrowhead), and its apparent correlation with the extent of *MORF2* suppression, suggests that this band represents a precursor *ndhB* transcript retaining a 685-bp Group II intron (TAIR, https://www.arabidopsis.org). To test this hypothesis, we included rigorous DNase treatment during RNA isolation and prior to cDNA synthesis, and validated the absence of genomic DNA contamination by demonstrating no RT-PCR amplification of the intron-containing nuclear gene *ACTIN2* in any of the 12 cDNA samples (Fig. 2b). These results support the conclusion that the larger *ndhB* amplicons originated from unspliced premature RNAs rather than from genomic DNA.

To determine the nature of these long-insert transcripts, we developed a Python script to extract reads containing insertions ranging from 650 to 700 bp from all three replicates of each genotype (Fig. 1; Method S3). Using BLASTN pairwise sequence alignment (Fig. 1; Method S4), we revealed that these inserts shared 98.9 ± 2.1% sequence identities and an average alignment length of 685 ± 5 bp with the reference *ndhB* Group II intron (Table S3), indicating that they are indeed unspliced intron sequences present in the premature *ndhB* transcripts.

We next quantified the frequency of these unprocessed *ndhB* transcripts in each genotype. Both *P1-10* and *P1-12 iCRISPRi-MORF2* lines treated with 5 μM Dex exhibited statistically higher frequencies of premature *ndhB* transcripts compared to WT (Table 3), consistent with the more intense high-molecular-weight bands observed in their RT-PCR products (Fig. 2b). While the *KRAB* control line also showed a slightly elevated mean frequency compared to WT, this difference was not statistically significant (Table 3).

## Discussion

Despite the well-recognized biological importance of RNA editing in plant organelles, quantifying editing efficiencies with high accuracy and scalability remains technically challenging (Tang *et al*., 2024). Traditional Sanger sequencing approaches, while simple and accessible, lack the resolution and reproducibility necessary for quantitative comparisons across multiple sites and conditions. Meanwhile, advanced next-generation sequencing (NGS) strategies, such as targeted amplicon-seq and rRNA-depleted total RNA-seq, require complex workflows, substantial sequencing depth, and high costs, making them impractical for routine applications, particularly for targeted gene analysis or research conducted without access to extensive sequencing infrastructure.

To address these methodological limitations, we developed a rapid, cost-effective, and reproducible long-read sequencing approach, termed premium PCR sequencing, performed by Plasmidsaurus using Oxford Nanopore Technology, for digital quantification of RNA editing and transcript maturation at targeted organellar genes. By combining barcoded high-fidelity PCR amplicons with nanopore sequencing and custom in-house bioinformatic pipelines, this method enables direct and absolute quantification of C-to-U editing events with single-molecule resolution across multiple editing sites. Importantly, the protocol avoids labor-intensive gel purification, fragmentation, or adapter ligation steps, thus offering significant time and cost advantages over conventional NGS methods.

Our application of premium PCR sequencing to *ndhB* and *ndhD*, two chloroplast transcripts containing numerous editing sites, demonstrated its robust read coverage, high reproducibility across biological replicates, and substantial cost savings. The editing efficiency data obtained with this method not only reproduced previous findings based on Sanger sequencing (Yapa *et al*., 2023), but also extended them by resolving quantitative differences in a MORF2 dosage-dependent manner (Figs. 2a and 3). Notably, we observed a statistically significant reduction in editing efficiency at six *ndhB* and two *ndhD* sites in the strongly repressed iCRISPRi line *P1-12*, while only modest or no changes were detected in the mildly repressed line *P1-10* and the *KRAB* control. This gradient of editing defects reinforces the central role of MORF2 as a dosage-sensitive regulator of plastid RNA editing. It highlights the capacity of our method to capture subtle yet functionally relevant differences. Additionally, our practice of barcoded multiplexing, which combines three replicates and two genes into one premium PCR sequencing reaction, reduces the cost to only $30 per reaction (i.e., $10 per replicate), dramatically lowering the cost compared to lncRNA-Seq, which typically averages around $300 per replicate.

Beyond RNA editing, our approach also uncovered a striking accumulation of intron-retaining *ndhB* transcripts in Dex-treated *iCRISPRi-MORF2* lines. The increased frequency of reads containing 650–700 bp insertions, corresponding to the unspliced group II intron, directly correlated with the extent of MORF2 suppression (Fig. 2a and Table 2). These results not only validate the presence of intron-containing isoforms observed by gel electrophoresis (Fig. 2b), but also provide direct molecular evidence supporting the proposed role of MORF2 in coordinating RNA maturation and splicing. The observed intron retention phenotype upon MORF2 reduction in Dex-treated *iCRISPRi-MORF2* lines expands its known molecular function in chloroplast RNA processing, which is likely attributable to its holdase chaperon activity for broad involvement in chloroplast processes, including RNA editing (Takenaka *et al*., 2012), splicing (this study), tetrapyrrole biosynthesis (Yuan *et al*., 2022), and retrograde signaling (Yapa *et al*., 2023). This discovery of MORF2’s involvement in *ndhB* Group II intron splicing further underscores the utility of our platform in resolving transcript isoforms and splicing intermediates at the single-molecule level.

**Table 2.** Frequency of unspliced *ndhB* transcripts in four genotypes.

|  | **WT** |  |  | **KRAB** |  |  | **P1_10** |  |  | **P1-12** |  |  |
| --- | --- | --- | --- | --- | --- | --- | --- | --- | --- | --- | --- | --- |
| Replicates | R1 | R2 | R3 | R1 | R2 | R3 | R1 | R2 | R3 | R1 | R2 | R3 |
| Unspliced Reads | 4 | 4 | 4 | 10 | 15 | 5 | 21 | 8 | 4 | 40 | 27 | 9 |
| Aligned Reads | 531 | 439 | 196 | 614 | 471 | 215 | 585 | 337 | 186 | 501 | 330 | 122 |
| Frequency<br>(mean ± SD) | 1.2 ± 0.7 | | | 2.4 ± 0.8 | | | 2.7 ± 0.8* | | | 7.8 ± 0.4** | | |

† Frequency values represent the mean ± SD percentage of reads containing a 650–700 bp insert relative to total aligned reads per replicate. R1-R3 indicate three independent biological replicates as in Table 2.
\* P < 0.05; \*\*: P < 0.01 (Student's *t*-test, one-tailed, unequal variance)

A key innovation of this study lies in the suite of bioinformatic tools and scripts we developed to streamline every stage of the data processing workflow, from barcode demultiplexing and strand reorientation to pseudo-genome alignment and custom intron retention analysis (Fig. 1, Method S1-4). These bioinformatics tools, written in Perl and Python, are modular, lightweight, and optimized for full-length amplicon data from Oxford Nanopore platforms. By eliminating the need for large-scale RNA-seq pipelines and complex annotation-dependent software, our open-source scripts empower other laboratories to adopt long-read digital RNA editing and splicing variant analysis with minimal computational overhead. We anticipate that these resources will be broadly useful for studying plastid and mitochondrial RNA maturation, RNA-editing factors, and stress-responsive transcript isoform dynamics in a wide range of plant species.

In summary, our premium PCR sequencing and analysis platform represents a versatile and accessible approach for targeted analysis of RNA editing and processing in plant organelles. Its key strengths include simplicity, affordability, single-molecule precision, robust reproducibility, and scalable throughput across multiple genotypes or treatment conditions. By coupling a streamlined experimental workflow with open-access analytical tools, this method lowers the barrier to entry for plant biologists interested in RNA metabolism and supports a wide range of applications from basic research to translational crop biotechnology.

## Experimental Procedures

### Plant Materials, Growth, and Treatments

The iCRISPRi transgenic lines targeting *MORF2* were developed as described in our previous study (Yapa *et al*., 2023). Briefly, the dCas9-KRAB fusion protein was expressed under a Dex-inducible promoter (Aoyama and Chua, 1997), while the sgRNA targeting the TSS of *MORF2* was driven by the Arabidopsis *U6-26* promoter (Xing *et al*., 2014). Four genotypes were used in this study, including WT, *KRAB* control (expressing Dex-inducible *dCas9-KRAB* only), and two independent iCRISPRi lines (*P1-10* and *P1-12*). Seedlings were grown on half-strength Murashige and Skoog (1/2 MS) plates supplemented with 5 μM Dex under long-day photoperiod conditions (16 h light at 125 μmol m^-2^ sec^-1^, 21°C; 8 h dark, 19°C). Seven-day-old seedlings were harvested for RNA extraction.

### RNA Extraction, Reverse Transcription, and PCR Amplification

Total RNA was extracted using the NucleoSpin RNA Plus kit (Macherey-Nagel) according to the manufacturer’s protocol, followed by DNase I treatment (Thermo Fisher Scientific) to remove residual genomic DNA. For reverse transcription, 0.5 μg of oligo(dT)25 (IDT) and random hexamer primers (Thermo Fisher Scientific) were used in reactions with SuperScript III (Thermo Fisher Scientific) to capture full-length transcripts of *ndhB* and *ndhD*. PCR amplification was performed in a total volume of 50 μL using gene-specific primers flanking the full-length coding regions of *ndhB* and *ndhD*, with Phusion High-Fidelity DNA Polymerase (New England Biolabs) under standard cycling conditions. PCR amplicons were visualized on 1% agarose gels stained with ethidium bromide and quantified using a Qubit Flex Fluorometer with the dsDNA Broad Range (BR) assay kit (Thermo Fisher Scientific).

### PCR Amplicon Pooling and Nanopore Sequencing

Forward primers for *ndhB* and *ndhD* PCR products were barcoded for each of the three biological replicates (Tables S2 and S4). PCR products were cleaned using the NucleoSpin Gel and PCR Clean-up Kit (Macherey-Nagel) and pooled by genotype. Equimolar mixtures of *ndhB* and *ndhD* amplicons were prepared based on DNA concentrations and product lengths to ensure unbiased representation in the sequencing library (Table S1). Each pooled sample (four total) was prepared for nanopore sequencing using the Q20+ Kit V14 chemistry (Oxford Nanopore Technologies) and sequenced on a MinION flow cell (R10.4.1 chemistry) through a commercial service (Plasmidsaurus).

### Raw Data Processing and Barcode Demultiplexing

Raw FASTQ reads were basecalled and demultiplexed using Guppy (Plasmidsaurus). To ensure consistent strand orientation, a custom Perl script was developed using *Bio::SeqIO*, *Bio::Seq*, and *Bio::Seq::Quality* modules to reorient each read to the sense strand of *ndhB* and *ndhD* (Method S1). Based on unique 8-nt barcodes in the forward PCR primers, clean FASTQ files representing three biological replicates were reconstructed for each genotype (12 files total).

### Alignment to Pseudo-genome and C-to-U Editing Quantification

A custom pseudo-genome was constructed containing six synthetic chromosomes corresponding to the barcoded *ndhB* (chr1-3) and *ndhD* (chr4-6) sequences (Appendix S2). The 12 clean FASTQ files reconstructed as above were aligned to the pseudo-genome using *minimap2* (v2.24) with the following parameters for full-length cDNA amplicon alignment: *-ax map-ont* (Method S2). The resulting SAM files were converted to coordinate-sorted BAM files using *samtools*. The Integrated Genomics Viewer (IGV, v2.16.0) was used to visualize alignments for manually counting cytidine (C) and thymidine (T) nucleotides at known RNA-editing sites. Editing efficiency was calculated as *T/(C+T) x 100%* for each site.

### Insert Size Profiling and Unspliced Transcript Quantification

To identify unprocessed *ndhB* transcripts retaining the Group II intron, a Python script was developed to parse CIGAR strings from aligned BAM files for extracting insertions with lengths from 650 to 700 bp (Method S3). Reads containing such large insertions in chr1, chr2, or chr3 (representing three barcoded *ndhB* sequences in the pseudo-genome) were counted and normalized to the total number of reads mapped to the respective chromosome. The frequency of unspliced *ndhB* transcripts was defined as the proportion of total reads with a 650–700 bp insertion.

### Extraction and Local Sequence Alignment of Inserted Introns

Reads containing large insertions were cross-referenced to the raw FASTQ files to retrieve their full insert sequences, which were saved into separate FASTA files grouped by genotype. A Perl pipeline was used to conduct local sequence alignments between these insert sequences and the *ndhB* Group II intron sequence using *BLASTN* with default parameters (Method S4). The sequence similarities and alignment length were extracted and analyzed.

### Statistical Analysis

Editing efficiencies and intron retention frequencies were compared among genotypes using Student’s *t*-test. Statistical significance was defined as *p* < 0.05. For multiple replicates, means and standard deviations (SD) are reported.

## Acknowledgments

This work was supported in part by an NSF CAREER award (MCB-1750361 to ZH).

## Data Availability Statement

The data supporting the findings of this study are included in the supplementary materials accompanying this article. The complete bioinformatics pipeline—including sequence data (Appendices S1–S3) and analysis scripts (Methods S1–S4)—is also available on GitHub: https://github.com/hua-lab/Organelle_RNA_sequencing_analysis.

## Author Contributions

Z.H. conceived the project, designed the research, performed the experiments, analyzed data, and wrote the paper.

## Supporting Information

**Table S1** Equimolar mixture of *ndhB* and *ndhD* PCR products included in the final pool for premium PCR sequencing.

**Table S2** Sequence depth of *ndhB* and *ndhD* from 12 barcoded PCR amplicons aligned to each gene-specific reference sequence across four genotypes.

**Table S3** BLASTN alignment statistics comparing unspliced intron sequences to the reference *ndhB* Group II intron.

**Table S4** Primers used in this study.

**Method S1.** Perl script “filter_barcoded_fastq_seqs.pl” for converting raw FASTQ reads into a strand-corrected format.

**Method S2.** Bash script “run_minimap_alignments.sh” for aligning strand-corrected FASTQ files to the custom pseudo-genome using minimap2.

**Method S3.** Python script “count_groupII_inserts_and_extract.py” for identifying and extracting intron-containing *ndhB* transcripts.

**Method S4.** Perl script “local_align_inserts_vs_intron_blast.pl” for BLASTN alignment of intron-containing *ndhB* transcripts against the reference *ndhB* Group II intron sequence.

**Appendix S1** Raw FASTQ files generated from premium PCR nanopore long-read sequencing.

**Appendix S2** Sequence of the synthetic pseudo-genome used for read alignment

**Appendix S3** Reference sequence of the Group II intron from the *ndhB* gene

