## Supplementary material for "Rapid and Cost-Effective Digital Quantification of RNA Editing and Maturation in Organelle Transcripts": Table S4

**Table S4.** Primers used in this study.

| **Reactions** | **Primer Name** | **Sequence** |
| --- | --- | --- |
| Multiplexing RT-PCR | Rep1_*ndhB*_For | **ATGCTAGC**TTTTTGGCCTAATTCTTCTTCTGA |
|  | Rep2_*ndhB*_For | **CGTACGTA**TTTTTGGCCTAATTCTTCTTCTGA |
|  | Rep3_*ndhB*_For | **TACGATCG**TTTTTGGCCTAATTCTTCTTCTGA |
|  | *ndhB*_Rev | AATCGCAATAATCGGGTTCATT |
|  | Rep1_*ndhD*_For | **ATGCTAGC**AACAACTCGAAGTATGGGTC |
|  | Rep2_*ndhD*_For | **CGTACGTA**AACAACTCGAAGTATGGGTC |
|  | Rep3_*ndhD*_For | **TACGATCG**AACAACTCGAAGTATGGGTC |
|  | *ndhD*_Rev | GGCAATGCAAGGGAAGCC |
| qPCR | *MORF2*_qPCR_For | AGCTAAGCCAGTATGAGGATGA |
|  | *MORF2*_qPCR_Rev | CCAAAGTAAGCCCGGTCCAT |
|  | *ACT2*_qPCR_For | GGCATCACACTTTCTACAATGAG |
|  | *ACT2*_qPCR_Rev | ACCCTCGTAGATTGGCACAG |
|  | *PP2A*_qPCR_For | AAGCTTGGTGCTCTTTGCAT |
|  | *PP2A*_qPCR_Rev | CCATTACTGGAGCGAGAAGC |

† The underlined and bolded 8-nucleotide sequences at the 5′ end of the forward primers for *ndhB* and *ndhD* indicate the barcodes used for multiplexed RT-PCR across three biological replicates.
