## Supplementary material for "Rapid and Cost-Effective Digital Quantification of RNA Editing and Maturation in Organelle Transcripts": Table S3

|  | **WT** | **KRAB** | **P1_10** | **P1_12** | **ALL** |
| --- | --- | --- | --- | --- | --- |
| Sequence similarity (%; mean ± sd) | 99.1 ± 0.8 | 98.8 ± 2.4 | 99.1 ± 1.5 | 98.8 ± 2.0 | 98.9 ± 1.9 |
| Alignment length (nt; mean ± sd) | 686 ± 2 | 685 ± 5 | 685 ± 4 | 684 ± 6 | 685 ± 5 |

**Table S3.** BLASTN alignment statistics comparing unspliced intron sequences to the reference *ndhB* Group II intron.
