## Supplementary material for "Rapid and Cost-Effective Digital Quantification of RNA Editing and Maturation in Organelle Transcripts": Table S2

| **Sample** | **Barcode** | ***ndhB*** | ***ndhB* reads** | ***ndhD*** | ***ndhD* reads** |
| --- | --- | --- | --- | --- | --- |
| WT (Replicate 1) | ATGCTAGC | chr1 | 531 | chr4 | 636 |
| *KRAB* (Replicate 1) | ATGCTAGC | chr1 | 614 | chr4 | 535 |
| *P1_12* (Replicate 1) | ATGCTAGC | chr1 | 501 | chr4 | 470 |
| *P1_10* (Replicate 1) | ATGCTAGC | chr1 | 585 | chr4 | 594 |
| WT (Replicate 2) | CGTACGTA | chr2 | 439 | chr5 | 318 |
| *KRAB* (Replicate 2) | CGTACGTA | chr2 | 471 | chr5 | 365 |
| *P1_12* (Replicate 2) | CGTACGTA | chr2 | 330 | chr5 | 251 |
| *P1_10* (Replicate 2) | CGTACGTA | chr2 | 337 | chr5 | 341 |
| WT (Replicate 3) | TACGATCG | chr3 | 196 | chr6 | 258 |
| *KRAB* (Replicate 3) | TACGATCG | chr3 | 215 | chr6 | 271 |
| *P1_12* (Replicate 3) | TACGATCG | chr3 | 122 | chr6 | 170 |
| *P1_10* (Replicate 3) | TACGATCG | chr3 | 186 | chr6 | 255 |

**Table S2**. Sequence depth of *ndhB* and *ndhD* from 12 barcoded PCR amplicons aligned to each gene-specific reference sequence across four genotypes.

† The barcoded *ndhB* and *ndhD* reference sequences correspond to chr1-3 and chr4-6, respectively, as organized in a custom pseudo-genome (Appendix S2).
