## Supplementary material for "Rapid and Cost-Effective Digital Quantification of RNA Editing and Maturation in Organelle Transcripts": Table S1

**Table S1**. Equimolar mixture of *ndhB* and *ndhD* PCR products included in the final pool for premium PCR sequencing.

| **Genotypes** | ***ndhB*** | | | | | ***ndhD*** | | | | |
| --- | --- | --- | --- | --- | --- | --- | --- | --- | --- | --- |
|  | Length(bp) | Conc (ng/μL) | Mix Volume (μL) | Final  Conc (μM) | Reads | Length(bp) | Conc (ng/μL) | Mix Volume (μL) | Final  Conc (μM) | Reads |
| **WT** | 1405 | 96.0 | 5.2 | 0.041 | 1166 | 1355 | 60.0 | 8.0 | 0.041 | 1212 |
| ***KRAB*** | 1405 | 70.4 | 7.0 | 0.035 | 1300 | 1355 | 59.6 | 8.0 | 0.035 | 1171 |
| ***P1-12*** | 1405 | 87.6 | 8.9 | 0.050 | 953 | 1355 | 94.4 | 8.0 | 0.050 | 891 |
| ***P1-10*** | 1405 | 118 | 7.2 | 0.060 | 1108 | 1355 | 102.0 | 8.0 | 0.060 | 1190 |

† The table shows the initial concentrations and volumes used to prepare equimolar mixtures of *ndhB* and *ndhD* PCR products from each genotype.
