## Supplementary material for "Rapid and Cost-Effective Digital Quantification of RNA Editing and Maturation in Organelle Transcripts": Method S4

#!/usr/bin/env perl

# Script: local_align_inserts_vs_intron_blast.pl
# Tested on: macOS (ARM64) with Perl 5.40.1 and NCBI BLAST+ v2.16.0

# Description:
#   Performs local sequence alignment of unspliced ndhB insert reads (650–700 bp) 
#   against the reference ndhB Group II intron sequence using BLASTN.
#   Extracts alignment identity and length for each read and summarizes the results 
#   by genotype in a tab-delimited output table.

# Usage:
#   1. Save this script as "local_align_inserts_vs_intron_blast.pl" in the same directory as the BAM files (Method S2).
#   2. Ensure the "merged_fastas/" directory from Method S3 contains per-genotype merged FASTA files 
#      (e.g., WT_inserts.fasta, Krab_inserts.fasta).
#   3. Provide the reference intron FASTA file (e.g., "ndhB_intron.fasta").
#   4. Run the script:
#        perl local_align_inserts_vs_intron_blast.pl

# Output:
#   - A BLAST database is generated from "ndhB_intron.fasta"
#   - Each merged FASTA file is locally aligned against this database
#   - Output summary file: "ndhB_insert_vs_intron_blast.tsv", with columns:
#       Genotype | Read_ID | Percent_Identity | Alignment_Length

# Dependencies:
#   - NCBI BLAST+ (makeblastdb, blastn)
#   - Perl ≥ 5.10

# Citation:
#   Hua, Z. et al. (2025). Rapid and Cost-Effective Digital Quantification of RNA Editing and 
#   Maturation in Organelle Transcripts. *The Plant Journal*, [in review].

# Contact:


use strict;
use warnings;
use File::Basename;

# === Inputs ===
my $intron_fasta = "ndhB_intron.fasta";
my $merged_dir   = "merged_fastas";
my $output_tsv   = "ndhB_insert_vs_intron_blast.tsv";

# === Prepare BLAST database ===
print "Building BLAST database from $intron_fasta...\n";
system("makeblastdb -in $intron_fasta -dbtype nucl -out intron_db") == 0
    or die "Failed to make BLAST database.\n";

# === Open output file ===
open(my $out, '>', $output_tsv) or die "Cannot open $output_tsv: $!";
print $out join("\t", qw(Genotype Read_ID Percent_Identity Alignment_Length)), "\n";

# === Process each merged fasta file ===
opendir(my $dh, $merged_dir) or die "Cannot open $merged_dir: $!";
my @files = grep { /\.fasta$/ } readdir($dh);
closedir($dh);

foreach my $fasta (@files) {
    my $fasta_path = "$merged_dir/$fasta";
    my $genotype = $fasta;
    $genotype =~ s/_inserts\.fasta//;

    print "Running BLASTN for $genotype...\n";

    # Run blastn on the current file
    my $blast_out = "temp_blast.tsv";
    system("blastn -query $fasta_path -db intron_db -outfmt '6 qseqid pident length' -out $blast_out") == 0
        or die "Failed to run blastn for $fasta\n";

    # Parse results
    open(my $blast_fh, '<', $blast_out) or die "Cannot read BLAST output.\n";
    while (my $line = <$blast_fh>) {
        chomp $line;
        my ($read_id, $identity, $aln_len) = split(/\t/, $line);
        print $out join("\t", $genotype, $read_id, $identity, $aln_len), "\n";
    }
    close($blast_fh);
    unlink $blast_out;
}

close($out);
print "All done. Results saved in $output_tsv\n";
