## Supplementary material for "Rapid and Cost-Effective Digital Quantification of RNA Editing and Maturation in Organelle Transcripts": Method S3

#!/usr/bin/env python3

# Script: count_groupII_inserts_and_extract.py  
# Tested on: macOS (ARM64) with Python 3.11, Biopython v1.81, and pandas v2.2.2  
#
# Description:  
#   Identifies reads containing large insertions (650–700 bp) from sorted BAM files aligned to  
#   barcode-specific ndhB pseudo-genome chromosomes (chr1–3). Extracts corresponding unspliced  
#   reads from strand-corrected FASTQ files, saves them as individual and merged FASTA files,  
#   and outputs a summary table of intron-retention frequencies by genotype and barcode.  
#
# Usage:  
#   1. Save this script in the same directory as the 12 sorted BAM files.  
#   2. Ensure the strand-corrected FASTQ files are located in the parent directory (../)  
#      and include both barcode and genotype identifiers in their filenames.  
#   3. Set up and activate a virtual environment:  
#        python3 -m venv venv  
#        source venv/bin/activate  
#        pip install biopython pandas  
#   4. Run the script:  
#        python count_groupII_inserts_and_extract.py  
#   5. Deactivate the virtual environment when done:  
#        deactivate  
#
# Output:  
#   - ndhB_groupII_insertion_count_chr1-3.tsv: Summary table of unspliced read frequencies  
#   - insert_fastas/: Folder containing per-sample FASTA files with 650–700 bp insert reads  
#   - merged_fastas/: Folder containing genotype-wise merged FASTA files for alignment or BLASTN  
#
# Dependencies:  
#   - Python ≥ 3.6  
#   - Biopython ≥ 1.81  
#   - pandas ≥ 2.2.2  
#   - samtools (must be in your system PATH)  
#
# Citation:  
#   Hua, Z. et al. (2025). Rapid and Cost-Effective Digital Quantification of RNA Editing and  
#   Maturation in Organelle Transcripts. *The Plant Journal*, [in review].  
#
# Contact:  


import os
import re
import pandas as pd
import subprocess
from Bio import SeqIO
from glob import glob

# --- Configuration ---
barcode_chr_map = {
    "ATGCTAGC": ["chr1"],
    "CGTACGTA": ["chr2"],
    "TACGATCG": ["chr3"]
}
expected_genotypes = ["WT", "Krab", "10-1", "12-1"]
fastq_dir = "../"

# --- Setup output directories ---
os.makedirs("insert_fastas", exist_ok=True)
os.makedirs("merged_fastas", exist_ok=True)

# --- Genotype name parser ---
def parse_genotype(bam_name):
    for gt in expected_genotypes:
        if f"_{gt}" in bam_name:
            return gt
    return "Unknown"

# --- Process each BAM file ---
bam_files = [f for f in os.listdir() if f.endswith(".sorted.bam")]
groupII_results = []

for bam in bam_files:
    for barcode, chroms in barcode_chr_map.items():
        if barcode in bam:
            genotype = parse_genotype(bam)
            for chrom in chroms:
                try:
                    sam_output = subprocess.run(
                        ["samtools", "view", bam, chrom],
                        capture_output=True, text=True, check=True
                    )
                    lines = sam_output.stdout.strip().split("\n")
                    total_reads = len(lines)
                    cigar_lines = [(line.split('\t')[0], line.split('\t')[5]) for line in lines if len(line.split('\t')) > 5]

                    insert_ids = []
                    for read_id, cigar in cigar_lines:
                        insertions = list(map(int, re.findall(r'(\d+)I', cigar)))
                        if any(650 <= i <= 700 for i in insertions):
                            insert_ids.append(read_id)

                    count = len(insert_ids)
                    freq = round(count / total_reads, 5) if total_reads > 0 else 0.0

                    groupII_results.append({
                        "BAM File": bam,
                        "Barcode": barcode,
                        "Chromosome": chrom,
                        "Genotype": genotype,
                        "Unspliced ndhB-like Group II Insertions (650–700 bp)": count,
                        "Total Aligned Reads": total_reads,
                        "Frequency of unspliced ndhB transcripts": freq
                    })

                    # Match FASTQ file and write insert reads to FASTA
                    fastq_matches = [f for f in os.listdir(fastq_dir) if barcode in f and genotype in f and f.endswith(".fastq")]
                    if fastq_matches:
                        fastq_path = os.path.join(fastq_dir, fastq_matches[0])
                        fasta_out = f"insert_fastas/{genotype}_{barcode}_{chrom}_inserts.fasta"
                        with open(fasta_out, "w") as fasta_outfile:
                            count_written = 0
                            for record in SeqIO.parse(fastq_path, "fastq"):
                                if record.id in insert_ids:
                                    SeqIO.write(record, fasta_outfile, "fasta")
                                    count_written += 1
                            if count_written == 0:
                                os.remove(fasta_out)

                except subprocess.CalledProcessError as e:
                    print(f"[ERROR] samtools failed on {bam}, {chrom}: {e}")

# --- Save TSV summary ---
df = pd.DataFrame(groupII_results)
df.to_csv("ndhB_groupII_insertion_count_chr1-3.tsv", sep="\t", index=False)

# --- Merge per-genotype FASTA files ---
for gt in expected_genotypes:
    fasta_files = glob(f"insert_fastas/{gt}_*_inserts.fasta")
    merged_fasta_path = f"merged_fastas/{gt}_inserts.fasta"
    with open(merged_fasta_path, "w") as merged_fa:
        for f in fasta_files:
            for record in SeqIO.parse(f, "fasta"):
                SeqIO.write(record, merged_fa, "fasta")

print("TSV summary saved to 'ndhB_groupII_insertion_count_chr1-3.tsv'")
print("Merged FASTA files saved in 'merged_fastas/'")
