## Supplementary material for "Rapid and Cost-Effective Digital Quantification of RNA Editing and Maturation in Organelle Transcripts": Method S2

#!/bin/bash

# Script: run_minimap_alignments.sh
# Tested on: macOS (ARM64) with minimap2 v2.29 and samtools v1.21
#
# Description:
#   Aligns strand-corrected FASTQ files to a custom pseudogenome using minimap2,
#   followed by BAM file sorting and indexing using samtools.
#
# Usage:
#   1. Rename "Method S2.txt" as "run_minimap_alignments.sh".
#   2. Save this script in the same directory as:
#        - The strand-corrected FASTQ files
#        - The pseudo-genome file renamed as "pseudo_genome_ndhBD.fa"
#   3. Run the following commands:
#        chmod +x ./run_minimap_alignments.sh
#        ./run_minimap_alignments.sh
#
# Dependencies:
#   - minimap2
#   - samtools
#
# Citation:
#   Hua, Z. et al. (2025). Rapid and Cost-Effective Digital Quantification of RNA Editing and  
#   Maturation in Organelle Transcripts. *The Plant Journal*, [in review].
#
# Contact:


# Set the number of threads
THREADS=4

# Reference genome (pseudo-genome)
REF="pseudo_genome_ndhBD.fa"

# Make sure the reference is indexed for minimap2
if [ ! -f "$REF.mmi" ]; then
    echo "Indexing reference genome..."
    minimap2 -d "$REF.mmi" "$REF"
fi

# Create output directory
mkdir -p minimap_alignments

# List of filtered FASTQ files
FILES=(

TACGATCG_45CNXP_4_10-1.fastq
TACGATCG_45CNXP_3_12-1.fastq
TACGATCG_45CNXP_2_Krab.fastq
TACGATCG_45CNXP_1_WT.fastq
CGTACGTA_45CNXP_4_10-1.fastq
CGTACGTA_45CNXP_3_12-1.fastq
CGTACGTA_45CNXP_2_Krab.fastq
CGTACGTA_45CNXP_1_WT.fastq
ATGCTAGC_45CNXP_4_10-1.fastq
ATGCTAGC_45CNXP_3_12-1.fastq
ATGCTAGC_45CNXP_2_Krab.fastq
ATGCTAGC_45CNXP_1_WT.fastq

)

# Loop over each FASTQ file and run minimap2
for FILE in "${FILES[@]}"; do
    SAMPLE=$(basename "$FILE" .fastq)
    echo "Aligning $SAMPLE..."

    minimap2 -ax map-ont -t $THREADS "$REF.mmi" "$FILE" | \
        samtools sort -@ $THREADS -o minimap_alignments/${SAMPLE}.sorted.bam

    samtools index minimap_alignments/${SAMPLE}.sorted.bam
    echo "Alignment complete: minimap_alignments/${SAMPLE}.sorted.bam"
done

echo "All alignments completed and indexed."
